# The evolution of male weapons is associated with the type of breeding site in a clade of Neotropical frogs

**DOI:** 10.1101/2024.08.15.608145

**Authors:** Erika M. Santana, Daniel S. Caetano, Alexandre V. Palaoro, Glauco Machado

**Affiliations:** Universidade de Sao Paulo; Towson University; Universidade Federal do Parana

**Keywords:** evolutionary lability, hypertrophied limbs, male-male competition, sexual dimorphism, sexual selection, thumb spines

## Abstract

Male weaponry evolution is often linked to resource or female defense polygyny. This pattern emerges from highly studied taxa, such as arthropods and mammals. However, whether factors such as breeding site type influence weaponry evolution remains an open question. We explore this question using frogs of the subfamily Leptodactylinae, where males of species spawning in exposed sites (water bodies) fight to hold oviposition sites or dislodge rivals during amplexus. Conversely, males of species spawning in concealed sites (ground nests and crevices) rarely engage in intense physical contests. We hypothesize that male weaponry evolution would be associated with reproduction in exposed sites. Using two complementary phylogenetic comparative methods, we first found a macroevolutionary correlation between male weapons and breeding site type. The presence of weapons is consistently associated with exposed sites, while their absence is linked to concealed sites. We explore how this macroevolutionary pattern can also be influenced by female mate choice. Second, we found that both gains and *losses* of weapons occurred more frequently in species spawning in exposed sites. This unexpected finding suggests that the dynamics of weapon evolution are more intricate than anticipated. We propose some explanations aspiring to stimulate further investigations with other external fertilizer species.

## 1. Introduction

Females of many species use specific places or limited resources to lay their eggs. In certain taxa, the limited resources are nests, such as burrows or natural cavities, which can be economically defended by males against competitors (crustaceans: e.g., [1]; arachnids: e.g., [2]; insects: [3]; fish: e.g., [4]; anurans: e.g., [5]; birds: e.g., [6]). The monopolization of these oviposition sites involves male-male contests, which may drive the evolution of fighting-related traits used to threaten or physically coerce rivals [7]. In fact, the presence of fighting-related traits (hereafter ‘weapons’) seems to be more frequent in species showing resource defense polygyny than other types of mating systems, such as leks or scramble competition [7–9]. Another type of mating system in which weapons are common is female defense polygyny, in which males directly compete for access to females [7–9]. In both resource and female defense polygyny, males bearing larger or stronger weapons typically prevail in the contests [10,11], gaining access to a greater number of females and thus achieving a higher reproductive success compared to males bearing smaller or weaker weapons [12].

Despite the widespread occurrence of weapons in animals, empirical studies on weaponry evolution are concentrated in a few groups, such as crustaceans, beetles, and mammals [7,9,11]. A taxon for which the selective pressures underlying weaponry evolution are poorly understood are the order Anura (Amphibia). Anurans comprise a highly diverse clade of vertebrates in which male weaponry, including fangs, hypertrophied arms, and thumb and chest spines, evolved independently in several families [5,9,13]. In some of these families, weapons are present in species in which males build energetically costly nests that can be used by females as oviposition sites (e.g., [14–16]). In other families, weapons are present in species with exclusive paternal care in which males fight for the possession of oviposition sites that can be visited and used by multiple females (e.g., [17]). Weapons also occur in species in which males do not care for the offspring but defend suitable oviposition sites (e.g., [18–21]). Finally, there are species in which males do not defend reproductive sites, but instead use their weapons to gain access directly to females in mating aggregations (e.g., [22,23]). Thus, as in other taxa, weaponry evolution in anurans also appears to be associated with resource or female defense, which are mating systems characterized by frequent physical contests among males [5,9,13]. However, whether factors such as the *type of breeding site* may also influence weaponry evolution remains an open question.

Male-male contests have been reported for several frog species of the Neotropical subfamily Leptodactylinae (Leptodactylidae), in which females lay their eggs in exposed water bodies or concealed sites, such as small burrows excavated by males on the ground, or less frequently, within rock crevices defended by males ([5,13]; figure 1A-B). In species where females lay eggs in concealed sites, such as those belonging to the genera *Lithodytes*, *Adenomera*, and *Leptopdactylus* of the *fuscus* group, contests primarily involve aggressive calls. Additionally, males may signal aggression by tapping their limbs on the ground, striking each other with snouts and limbs, or leaping onto rivals. However, instances of male-male grasping have not been reported, and the presence of weaponry traits is rare (e.g., [24–27]). In contrast, species where females lay eggs in exposed sites, such as water bodies or flooded excavated basins, exhibit more frequent aggressive physical contests, accompanied by the possession of weapons (e.g., [5,28–32]). One such trait is the presence of *corneous thumb spines* (figure 1C), which are used to stab rivals [9,13]. For instance, in *Le. pentadactylus*, males engaged in amplexus may use their thumb spines to fend off approaching rivals [28]. Another weapon is *hypertrophied arms* (figure 1D), also used for grasping females during amplexus [5,33]. In species like *Le. labyrinthicus*, *Le. latrans*, and *Le. pentadactylus*, males employ their hypertrophied forearms to grasp each other in a belly-to-belly position during contests [28–31,34]. Additionally, males of certain species have *pectoral* or *chest spines* (figure 1D), which are pressed against the rival’s chest in the *Leptodactylus* species in which males aggressively grasp each other in a belly-to-belly position [34]. These chest spines may also serve to reduce the likelihood of an amplexed male being displaced by others when direct competition for females is intense [29]. Despite their potential alternative uses, chest spines are specifically employed during contests [34].

**Figure 1.**
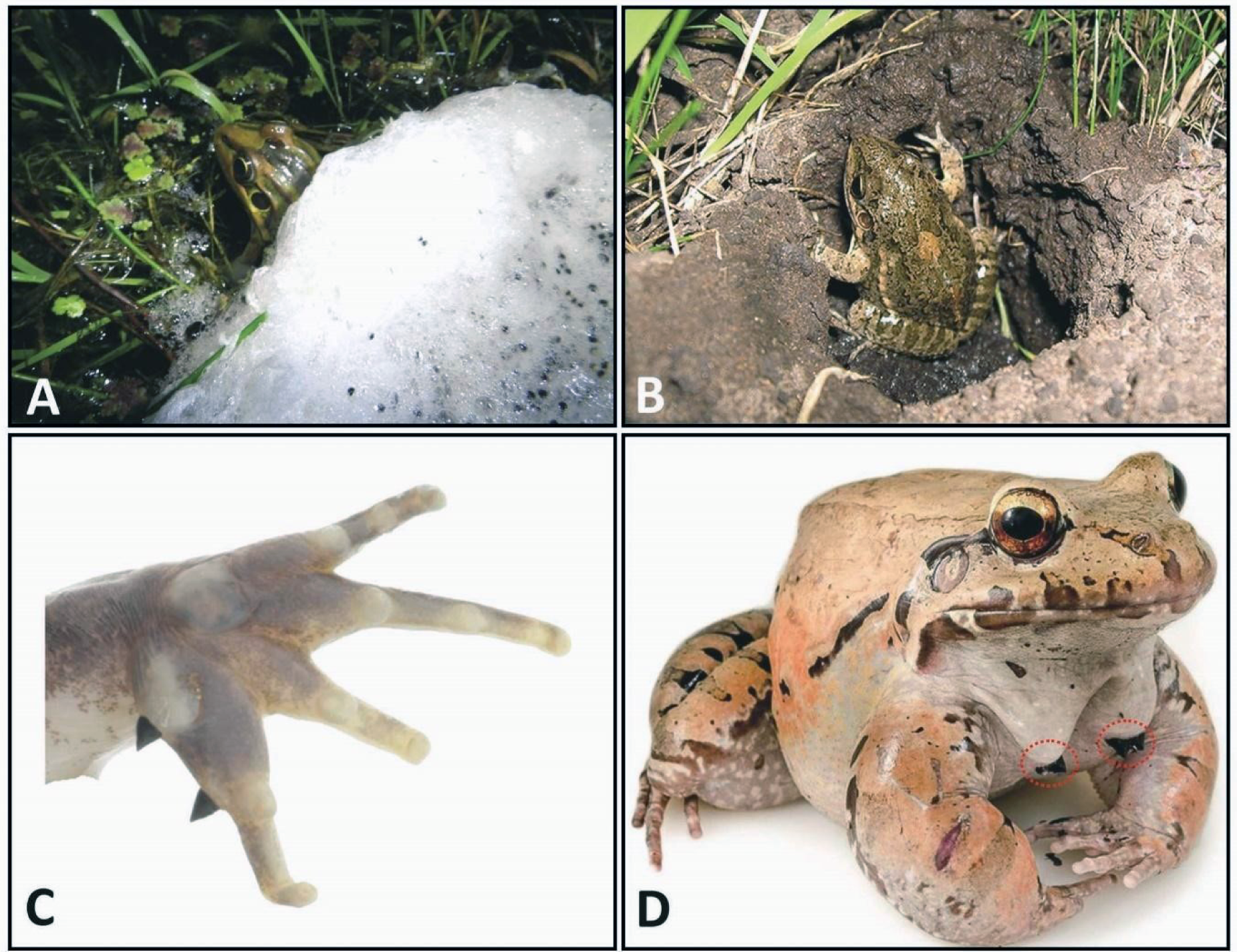
(A) Female of *Leptodoctylus latrans* inside an exposed foam nest on the water surface (photo by Florestas Pampeanas and Biofaces). (B) Male of *Le. latinasus* at the entrance of a concealed subterranean nest (photo by Raúl Maneyro). Note that the cap of the nest has been removed for the photograph to be taken. (C) Detail of the left forehand of a male *Le. validus* showing the thumb spines (modified from [41]). (D) Male of *Le. savagei* showing the hypertrophied forearms and the chest spines, which are the black, pointy projections emerging from the chest and surrounded by dashed red ellipses (photo by Felipe Villegas). The use of all images was authorized by the authors.

Based on scattered behavioral data, it appears that the intensity of male-male contests and the presence of weapons in Leptodactylinae frogs are associated with the type of breeding site. In species that spawn in exposed sites, two or more males can approach the same female before or during egg laying (e.g., [35]), and are expected to compete for the position closest to the female, as it likely increases their chances of fertilizing her gametes [36]. Additionally, females in these species select certain spawning sites [37], and males frequently engage in physical contests for the monopoly of these sites [29]. In both situations, being a successful competitor requires males to exclude rivals through physical contests, which may drive weaponry evolution [7,9,13]. In contrast, in species that spawn in concealed sites, males attract females by calling from inside burrows. Females choose these males by their call, enter the nest of the chosen male, and seal it after the eggs are laid (e.g., [38,39]). In these concealed sites, the number of males capable of entering and remaining close enough to release sperm onto female gametes is limited because the nest owner can effectively block the nest entrance with his body (e.g., [26,40]), which makes the access of rival males to the ovipositing female unfeasible. Consequently, males would not invest in aggressive interactions to steal fertilizations, which restricts weaponry evolution.

Following the rationale presented above, we hypothesized that reproduction in exposed breeding sites would correlate with the presence of male weapons over evolutionary time in Leptodactylinae frogs. Conversely, we hypothesized that concealed breeding sites would be associated with the absence of male weapons. Throughout the evolutionary history of this subfamily, transitions from reproduction in exposed breeding sites on the water surface to concealed breeding sites inside burrows have occurred multiple times [42]. Similarly, various weapons have been acquired and lost within the clade, providing a unique opportunity to investigate whether the evolution of male weapons is linked to the type of breeding site used by females. To test our hypotheses, we employed two complementary phylogenetic comparative approaches, and our findings demonstrate an evolutionary correlation between the presence of weapons and reproduction in exposed breeding sites. Furthermore, our results reveal that both gains and *losses* of weapons primarily occur in species that spawn in exposed sites, and at a higher frequency than in concealed sites. This unexpected outcome underscores the significance of breeding site exposure in the evolution of male weaponry and suggests that other factors may also influence weaponry evolution in Leptodactylinae frogs.

## 2. Methods

### (a) Study group

The subfamily Leptodactylinae is composed of 118 species currently divided into four genera [43]: *Adenomera* (30 species), *Hydrolaetare* (3 species), *Leptodactylus* (84 species), and *Lithodytes* (1 species). In all species of *Adenomera* and *Lithodytes*, females lay eggs in concealed breeding sites on land [28,44], whereas in the genus *Leptodactylus* females lay eggs either in exposed or concealed breeding sites ([42]; figures 1A-B). The breeding behavior of the three species of *Hydrolaetare* is still unknown.

### (b) Data collection

We conducted an extensive literature search for information on the type of breeding site and presence/absence of thumb spines, hypertrophied arms, and chest spines in males of each species of Leptodactylinae. We classified the breeding sites into two types: (1) *exposed*, including species with any kind of unprotected egg masses laid on the water surface or flooded excavated basins (reproductive modes 44-48, 50, 56, and 59 from [45]), and (2) *concealed*, including species with protected egg masses laid in concealed sites, such as inside terrestrial subterranean nests (reproductive modes 60, and 62-64 from [45]) or inside rock crevices (particular case of reproductive mode 48 from [45] when egg masses are isolated). We only included in our final dataset species that have all the information for any of the traits required (i.e., type of breeding site and the three morphological traits). Our final dataset includes 64 species: 1 of the genus *Lithodytes*, 4 of the genus *Adenomera*, and 59 of the genus *Leptodactylus* (Supplementary Information S1). Our dataset represents 54.2% of the total number of species currently recognized in the subfamily.

### (c) Phylogenetic tree

The most complete molecular tree published for the Leptodactylinae is from Sá et al. [46]. However, since then, other molecular studies with Leptodactylinae species have been published. Thus, we included new molecular data on *Adenomerea andreae* and *A. diptyx* [47], as well as on new species of the *Leptodcatylus latrans* complex [48–51], into the phylogeny originally proposed by Sá et al. [46]. We incorporated the newly obtained mitochondrial sequences (12S and 16S) and the nuclear Rhodopsin sequences from these recent molecular studies to the original alignment presented by Sá et al. [46] using a profile alignment in MAFFT [52]. This alignment approach retains the alignment of the previous phylogenetic tree while adding new sequences. We created two partitions, one with the mitochondrial and the other with the nuclear genes, and searched for the best-fitting model of molecular evolution for each partition using JModelTest2 [53]. Then, we used BEAST 2 [54], with a birth-death diversification model and a log-normal clock prior, to produce an ultrametric tree with node heights proportional to relative branching times of all Leptodactylinae species for which we found molecular data at GenBank (161 tips, see Supplementary Information S2). We ran 50 million generations of Markov Chain Monte Carlo and checked for convergence using Tracer [55].

Unfortunately, the molecular information for the species in Gazoni et al. [48], da Silva et al. [49], Carvalho et al. [50], and Magalhães et al. [51] were not available in GenBank, so we could not infer a larger tree based solely on molecular data. To circumvent this issue, we added 10 species following their putative sister species relationships along the midpoint of the branches (details at Supplementary Information S3). Finally, we computed the Maximum Clade Credibility tree (MCC) and pruned it to include only the 64 species of our dataset (Supplementary Information S1). The resulting tree is largely similar to Sá et al. [46], with only a few differences (compare figure 4 of Sá et al. [46] and figure 2 of this work), and was used for all subsequent comparative analyses.

**Figure 2.**
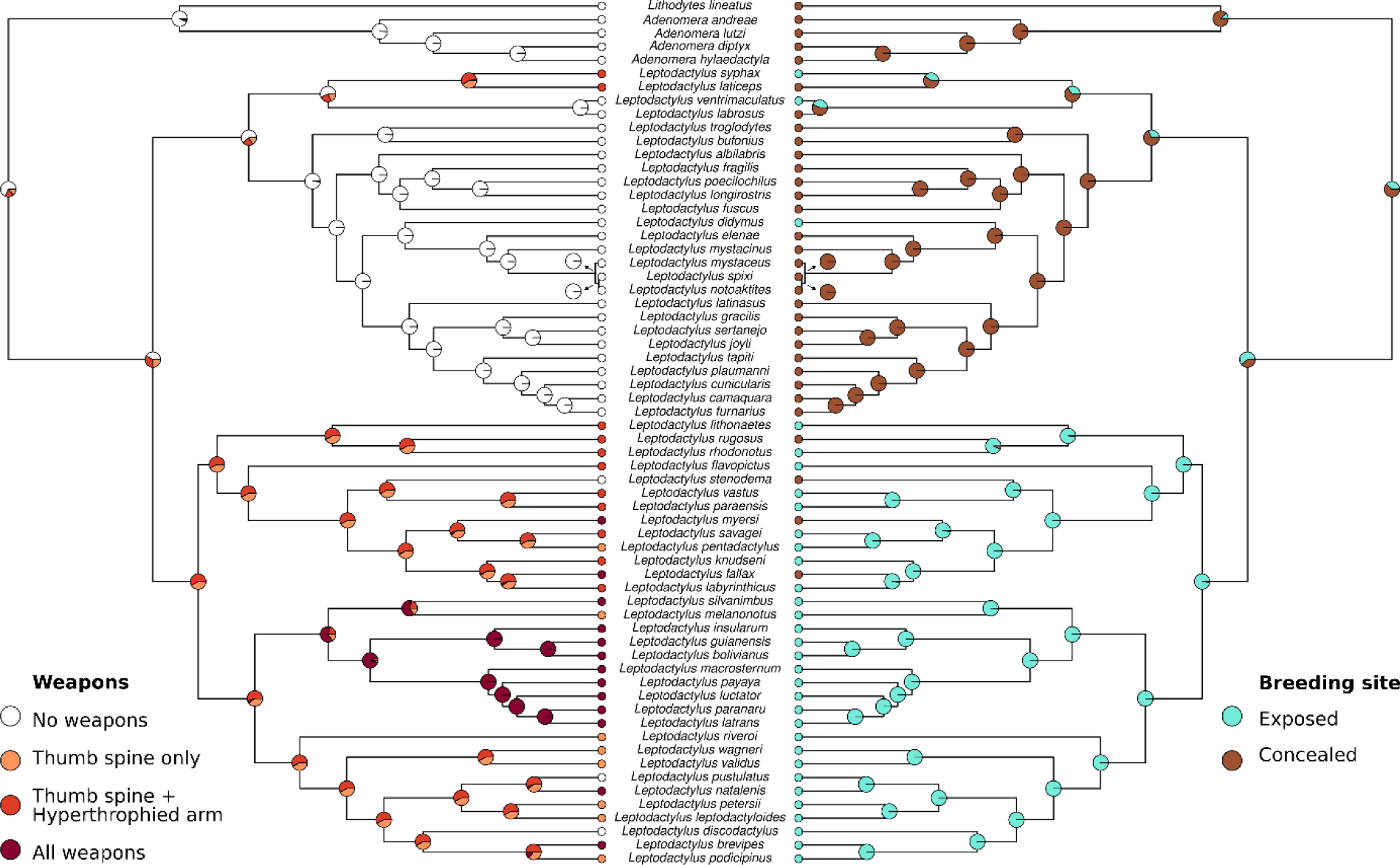
Reconstruction of the ancestral states of male weapons and breeding sites in Leptodactylinae frogs. Colors represent the four weapon states found in the group (weapons absent; only thumb spines present; thumb spines + hypertrophied arms present; thumb spines + hypertrophied arms + chest spines present) and the two states of breeding site (exposed or concealed). Ancestral states were reconstructed with a multi-state Markov model with transition rates freely varying but disallowing simultaneous transitions. Character histories were mapped for 1,000 iterations.

### (d) Comparative analyses

We used phylogenetic comparative methods to investigate the evolutionary association between male weapons (thumb spines, hypertrophied arms, and chest spines) and the type of breeding site (exposed and concealed). We employed two complementary approaches: first, we investigated the amount of evolutionary time during which the states of each trait (i.e., weapons and breeding site) co-occur along the branches of the phylogeny [56]; second, we investigated if the evolutionary transitions between the states of a trait (i.e., weapons) depend on the evolutionary transitions between the states of the other trait (i.e., breeding site) [57,58]. The first analysis allowed us to evaluate if the presence/absence of male weapons is associated with the type of breeding site throughout the evolution of the lineages. We quantified this by considering the amount of evolutionary time these traits co-occur in the phylogeny. The second analysis allowed us to investigate whether the likelihood of evolutionary gains and losses of male weapons differs between breeding sites. Specifically, this analysis investigated if gains of weapons are more likely in exposed breeding sites compared to concealed breeding sites.

For the first analysis, we estimated the co-occurrence of male weapons and type of breeding sites using independent Markov models with transition rates freely varying (the ARD model) but disallowing simultaneous transitions (i.e., only one state changes at any given time). Then, we simulated the history of male weapons and the type of breeding site using 1,000 stochastic mappings each and computed the co-occurrence of the presence/absence of male weapons with exposed or concealed breeding sites following Huelsenbeck’s Bayesian posterior predictive method, known as D-test [56]. The D-test measures the association between states along the branches of a tree (e.g., [59,60]). In our study, it means the proportion of evolutionary time male weapons co-occurred with each type of breeding site. To evaluate whether male weapons and breeding sites co-occur in the phylogeny more than expected if each trait evolved independently, we generated a null distribution of stochastic mappings by independently simulating the history of the evolution of male weapons and breeding sites. To do so, we used the transition rates estimated from the data and computed the likelihood of association as if these traits were truly independent.

We estimated the strength of the effect (measured in units of branch length) as the difference between the observed and null co-occurrence between states. Larger values indicate that the sum of branch lengths where the states co-occurred in the observed data was greater than in the null model. Negative values indicate that the sum of branch lengths where the observed states co-occurred in the observed data was lower than in the null model. According to our hypothesis, we expected a positive value when male weapons co-occur with exposed breeding sites and a negative value when male weapons co-occur with concealed breeding sites. Finally, we compared the degree of association in the observed data with the null distribution using a posterior predictive p-value [56].

For the second analysis, we applied Hidden-Rate Markov’s models of correlated evolution (hereafter HRMM) for discrete traits [57,58]. Since our study has a binary predictor (type of breeding site) and a four-state response trait (absence of weapons; only thumb spines; thumb spines + hypertrophied arms; thumb spines + hypertrophied arms + chest spines), we used a multi-state correlated model implemented in the R package *corHMM* [61]. To test for the evolutionary correlation between male weapons and the type of breeding site, we first estimated each of the transition models described below, both with and without the dependence of the type of breeding site. We used the Akaike Information Criterion corrected for sample size (AICc) to compare the fit of the dependent and independent models. We also included hidden state models to control for the potential effect of evolutionary heterogeneity among lineages in the results, thus avoiding issues with model inadequacy and type-I error [58,62–64]. A hidden state is considered an additional (binary) trait that, when it changes its state, can accelerate or decelerate the transition rates of the focal traits. Hence, models with hidden states can have multiple evolutionary rates along the phylogeny. We did not perform a HRMM test with a two-state response trait (presence and absence of weapons, collapsing all weapons into a single state) because doing so drastically reduced the number of transitions, thus decreasing the test’s power to detect any patterns.

To estimate the transitions between the four weapon states, we built the following transitions matrices: (1) all transitions between weapon states (absence and the three possibilities of presence, see above) can occur with distinct rates (hereafter ‘free model’); (2) all gains of weapons (transitions from absence to any of three states of presence) occur at the same rate, while losses occur at a different rate from gains, but at the same rate regardless of which trait is losing (hereafter ‘gains model’); (3) all transitions between the four weapon states (absence and three states of presence) happen under the same rate, i.e., rates of gains and losses did not differ (hereafter ‘equal model’). For this analysis, we expected that the concurrent models in which weapon evolution depends on the type of breeding site presented the best fit when compared with independent models (i.e., AICc < 2.0). Specifically, we expected that gains of weapons (i.e., rates of change from weapons absent to any of the three states in which weapons are present) would be more frequent when species spawn in exposed breeding sites. Additionally, we expected that losses of weapons (i.e., rates of change to weapons absent from any other state) would be more frequent when species spawn in concealed breeding sites.

By combining these two analyses, the D-test and HRMM, we can achieve a more comprehensive understanding of how male weapons and type of breeding site are evolutionarily linked. Specifically, the D-test estimates the proportion of the evolutionary time that each of the four weapon states co-occur with each of two types of breeding sites. According to the D-test, traits are considered correlated if they co-occur across the branches of the phylogeny more often than expected under the null model. This null model is produced by stochastic mapping simulations using parameters estimated from the observed data, ensuring that the null and alternative models have the same complexity (more details in Supplementary Material S3). In contrast, the test of evolutionary correlation, first proposed by Pagel [57] and later extended to include hidden states [58], focuses on transition rates among states and compares a model of dependent evolution (e.g., transitions between weapon states depend on the type of breeding site) with a model of independent evolution of traits. Thus, the D-test and HRMM explore distinct expectations of evolutionary correlation between discrete states.

## 3. Results

### (a) Weapons are mostly present in a single clade

Among the 64 Leptodactylinae species included in our dataset, 28 have at least one weapon. We found the following combinations of weapon traits: (a) only thumb spines (8 species), (b) thumb spines + hypertrophied arms (13 species), and (c) thumb spines + hypertrophied arms + chest spines (11 species). Regarding the type of breeding site, we found 32 species that spawn in exposed sites. Four of these species do not have any weapon, 8 have only thumb spines, 11 have thumb spines + hypertrophied arms, and 9 have thumb spines + hypertrophied arms + chest spines. The other 32 species spawn in concealed sites. Among them, 28 do not bear any weapon, 2 have thumb spines + hypertrophied arms, and 2 have thumb spines + hypertrophied arms + chest spines (Supplementary Information S1). The ancestral state for the Leptodactylinae clade is likely to be reproduction in concealed breeding sites and absence of male weapons (figure 2).

### (b) Weapons are associated with breeding in exposed sites

The D-test showed a strong relationship between exposed breeding sites and the presence of male weapons through evolutionary time (figure 2). The *presence* of thumb spines alone was associated with *exposed* breeding sites during 36% of the evolutionary history of the clade. The *presence* of thumb spines + hypertrophied arms was associated with *exposed* breeding sites during 27% of the evolutionary history of the clade. Finally, the *presence* of thumb spines + hypertrophied arms + chest spines was associated with *exposed* breeding sites during 26% of the evolutionary history of the clade. Combining the results, the *presence* of at least one of the three weapons was associated with *exposed* breeding sites during 89% of the evolutionary history of the clade (table 1). These results are accompanied by significant associations between the *presence* of any weapon trait and exposed breeding sites (p < 0.01, table 1). Conversely, the *absence* of the three weapons investigated here was significantly associated (p < 0.01, table 1) only with *concealed* breeding sites, and this association occurred during 85% of the evolutionary history of the clade. In summary, male weapons in Leptodactylinae frogs are frequently associated with reproduction in exposed sites, while the absence of male weapons is frequently associated with reproduction in concealed sites, and these traits co-occur more than expected if they evolved independently.

**Table 1.** Results of posterior predictive test (D-test) of co-occurrence between male weapons and type of breeding site (exposed or concealed) in Leptodactylinae frogs. We compared the degree of association between the presence/absence of weapons and the type of breeding site observed in our data with the null distribution of stochastic mappings generated by independently simulating the history of the evolution of male weapons and breeding sites. Rows show (i) marginal probabilities (models’ comparison with the posterior predictive p-values); (ii) temporal association (proportion of evolutionary history) between the type of breeding site and each combination of weapon traits (0 = trait absent; TS = thumb spines; HA = hypertrophied arms; CS = chest spines); and (iii) effect strength. The orientation of the arrows indicates if the observed association was higher (up) or lower (down) than expected if traits (weapons and breeding site) had evolved independently, i.e., without correlation.

| Type of breeding site | Statistics | Trait combination |  |  |  |
| --- | --- | --- | --- | --- | --- |
|  |  | 0 + 0 + 0 | TS + 0 + 0 | TS + HA + 0 | TS + HA + CS |
| Exposed | p-value | > 0.9 | < 0.01 | < 0.01 | < 0.01 |
|  | Association | ↓ 0.12 | ↑ 0.36 | ↑ 0.27 | ↑ 0.26 |
|  | Effect strength | -0.183 | 0.079 | 0.056 | 0.048 |
| Concealed | p-value | < 0.01 | > 0.9 | > 0.9 | > 0.9 |
|  | Association | ↑ 0.85 | ↓ 0.04 | ↓ 0.06 | ↓ 0.06 |
|  | Effect strength | 0.183 | -0.079 | -0.056 | -0.048 |

*(c) Gains and losses of weapons depend on the type of breeding site*

### (c) Gains and losses of weapons depend on the type of breeding site

The test of dependent evolutionary transitions between each weapon trait and the type of breeding site showed support for the dependent equal model, with transitions dependent on the type of breeding site (table 2). This means that gains and losses of weapons occur at the similar rates, but these rates differ between breeding sites, being higher in exposed than in concealed sites (figure 3). Gaining or losing weapons is 28 times more likely to occur in exposed breeding sites than in concealed breeding sites (figure 3). Furthermore, evolutionary transitions in weapon traits were estimated to happen much more frequently than changes in the type of breeding site, which is concentrated in certain clades (figure 4).

**Table 2.** Models of correlated evolution between male weapons and the type of breeding site in Leptodactylinae frogs. We present model specification, number of free parameters, and description of each model. Each model was run both with and without a hidden state. Models are sorted in ascending order of the values of the delta Akaike Information Criterion corrected for small sample sizes (ΔAICc). ΔAICc was calculated using the lower AICc as the base and subtracted from the rest of the AICc.

| Model specification | Free parameters | Model description | $\Delta\text{AICc}$ |
| --- | --- | --- | --- |
| (A) Equal dependent | 4 | All rates equal and dependent on the type of breeding site | 0 |
| (B) Equal independent | 3 | All rates equal, but independent of type of breeding site | 12.1 |
| (C) Free dependent | 14 | All rates are different and dependent on the type of breeding site | 12.8 |
| (D) Gains dependent | 6 | Separated rates of gains and losses and dependent on type of breeding site | 14.6 |
| (E) Equal dependent + hidden state | 10 | Same as model (A), including a hidden state | 22.5 |
| (F) Equal independent + hidden state | 8 | Same as model (B), including a hidden state | 24.8 |
| (G) Free independent | 8 | All rates are different, but independent of the type of breeding site | 33.2 |
| (H) Gains independent | 4 | Separated rates of gains and losses, but independent of type of breeding site | 38.6 |
| (I) Gains dependent + hidden state | 14 | Same as model (D), including a hidden state | 141.7 |
| (J) Gains independent + hidden state | 10 | Same as model (H), including a hidden state | 152.0 |

**Figure 3.**
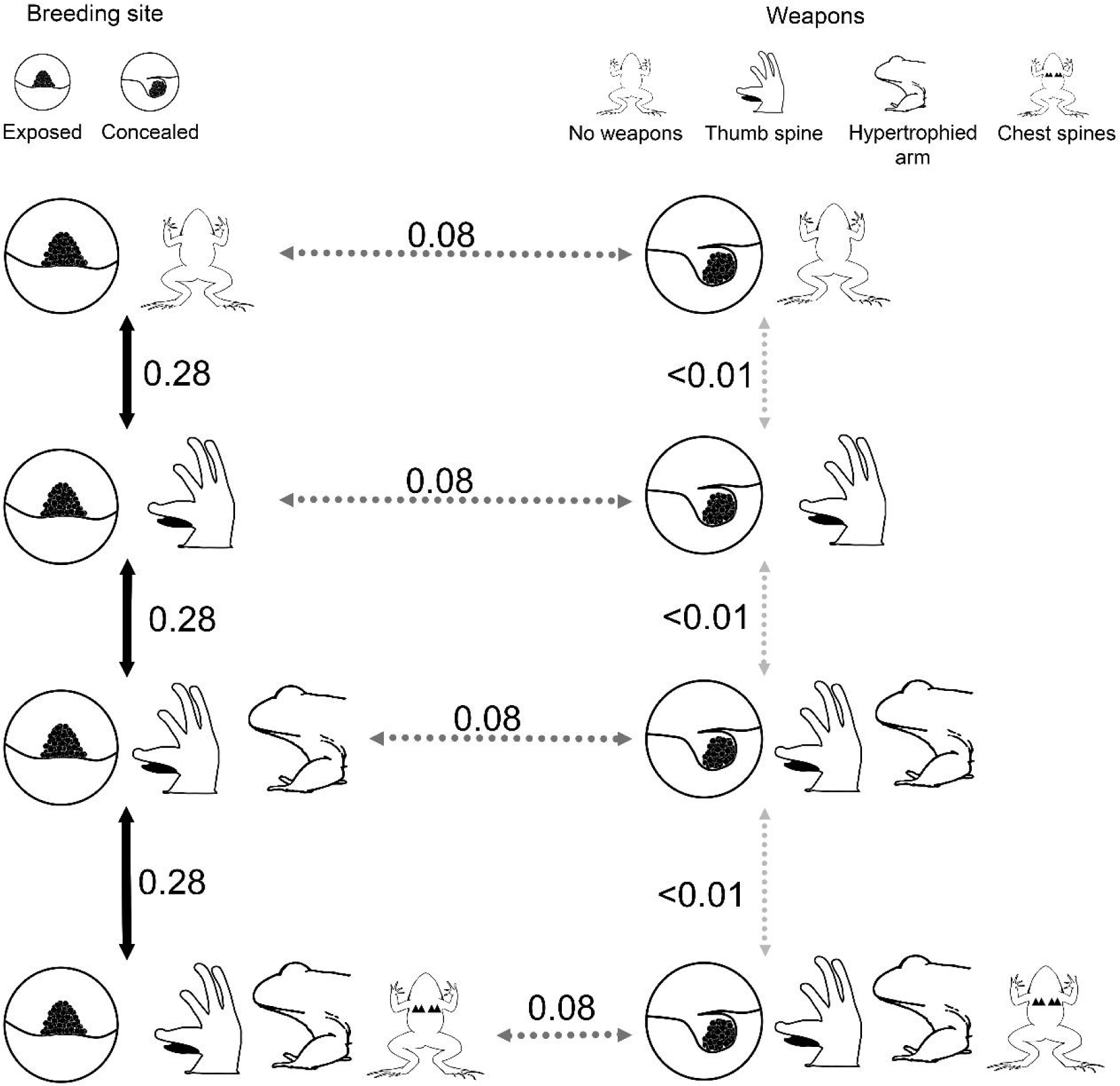
Evolutionary transition rates between the four weapon states (weapons absent; only thumb spines present; thumb spines + hypertrophied arms present; thumb spines + hypertrophied arms + chest spines present) and the two types of breeding site (exposed or concealed) in Leptodactylinae frogs. Estimates are made using the Hidden-Rate Markov [57,58] transition states method and the molecular tree of 64 species included in our dataset (figure 2). Arrows are weighted in color and shape by the evolutionary transition rates (numbers beside them): black solid arrows denote higher rates, gray dotted arrows denote lower rates, and pale gray dotted arrows denote the lowest possible estimated rate.

**Figure 4.**
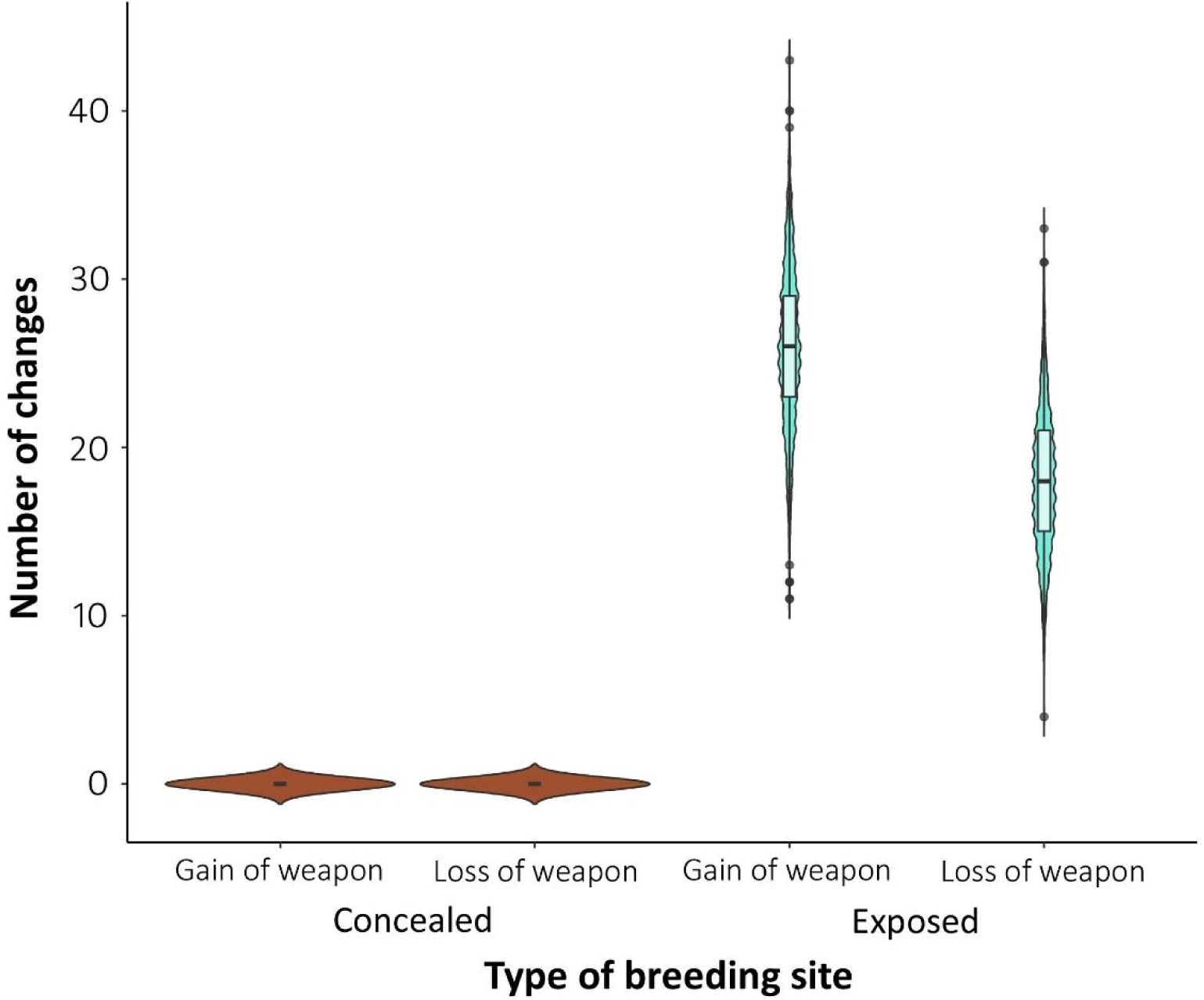
Number of gains and losses of male weapons in each type of breeding type calculated using the equal dependent model, which showed the best fit with our data (table 2). Males of Leptodactylinae frogs are losing and gaining weapons at a much higher rate in exposed breeding sites than in concealed breeding sites. Center line denotes the median, the box represents the quantiles and whiskers denote maximum and minimum values. Dots represent outliers (> 2 SD of the median), while the violin plot shows a density histogram of the data.

## 4. Discussion

While the evolution of weapons has been extensively documented and studied in taxa such as crustaceans [65], beetles [66], and mammals [67], it remains a relatively understudied subject in most taxa, especially those with external fertilization, including anurans. Our study addresses this knowledge gap by investigating whether the presence or absence of weapons correlates with the type of breeding site in a clade of Neotropical frogs. Scattered behavioral information indicates that Leptodactylinae species spawning in exposed sites exhibit a higher frequency of male-male aggression—either to fight for the possession of oviposition sites or to dislodge rival males during amplexus—compared to species spawning in concealed sites. Thus, we hypothesized that reproduction in exposed breeding sites would correlate with the presence of male weapons, while reproduction in concealed breeding sites would be associated with the absence of male weapons over evolutionary time. In our first comparative analysis, we found that the co-occurrence of weapons and reproduction in exposed sites spans 89% of the evolutionary history of the Leptodactylinae. Conversely, the absence of weapons was associated with reproduction in concealed sites, occurring during 85% of the evolutionary history of the clade. This association confirms our hypothesis that the presence of weapons and exposed breeding sites are evolutionarily correlated.

Theoretical and empirical studies on weaponry evolution typically correlate their presence with the type of mating system. Generally, weapons are more frequently found in species with female or resource defense mating systems [7–9]. This pattern have been confirmed in studies focusing on individual frog species [5,13,33], such as gladiator frogs from the family Hylidae and fanged frogs from the family Ranidae, in which males fiercely fight to protect nests or oviposition sites from usurpation by rivals [14,15,68]. However, among the Leptodactylinae species that construct nests on the ground, male agonistic interactions are almost entirely restricted to aggressive calls and occasional physical contact [24,26,40,69]. The absence of weapons in these species implies that nest-site defense alone cannot predict the presence of intense male-male fights and, consequently, cannot explain the patterns of weaponry evolution. This finding contrasts with a previous comparative study of dung beetles that found an evolutionary correlation between the presence of male horns and the defense of ground tunnels [70]. In fact, the absence of weapons in Leptodactylinae species that spawn inside concealed burrows reinforces our suggestion that the low risk of an amplectant male being dislodged by rivals in these sites reduces the probability or frequency of agonistic interactions, thereby decreasing the advantages associated with the presence of weapons. From a broader perspective, our comparative analysis indicates that the defense of concealed sites, such as burrows in the ground, does not necessarily lead to the evolution of male weaponry.

In frogs, the degree of female control over amplexus varies considerably among groups and can also be an important factor in predicting the evolution of male weaponry. In several species from the family Dendrobatidae, females visit the territory of multiple males before accepting one of them, and territorial males are never observed forcing amplexus (reviewed by [71]). Moreover, male-male agonistic interactions are primarily mediated through aggressive calls, and intense physical contact during contests is rare (e.g., [72–74]). Similar patterns are observed in Leptodactylinae species that spawn in concealed sites. For instance, in both *Leptodactylus bufonius* and *Le. fuscus*, males attract females to their nests by calling and never force amplexus. In these species, female acceptance seems to depend on a complex sequence of behaviors and courtship calls emitted by males and females, with the female only entering the nest voluntarily [75,76]. Thus, while the nest is crucial for males, having a nest does not guarantee female acceptance. The crucial role of female choice in defining male reproductive success likely attenuates the benefits of direct male-male competition for nest possession. This may further contribute to the low frequency of weapons in Leptodactylinae species that spawn inside concealed nests.

Contrary to the scenario described above, male forced amplexus appears to be common in anuran species that spawn in exposed sites, such as water bodies [5]. For instance, in the bufonid toad *Rhinella marina*, which spawns in exposed sites, male body size is negatively correlated with the dominant frequency of the call [77]. In phonotaxis experiments, females prefer male calls with low dominant frequency [78], indicating a preference for larger males [77]. However, the most successful males in achieving amplexus in the field are not the larger, but those with greater forelimb muscle mass [79]. These findings suggest that female mate choice is limited, and that the outcome of agonistic interactions is the main factor predicting male reproductive success. Moreover, it suggests that the evolution of weapons, such as hypertrophied arms, is related to direct male-male competition for females in exposed breeding sites. Although the evidence of forced amplexus is scarce for Leptodactylinae species that breed in exposed sites, we know that males attempt to dislodge other males during amplexus and that multiple males may be found competing for access to the same female [29,35]. When male reproductive success primarily depends on excluding rivals through physical contests, male weaponry is expected to evolve [7–9], and our results support this prediction.

Our second comparative analysis revealed that both gains and losses of weapons are 28 times more frequent in exposed breeding sites compared to concealed sites. We emphasize, however, that the results of the HRMM are sensitive to the number of transitions between trait states along the phylogeny [57,58]. Unfortunately, multiple transitions of the traits of interest may not occur along the evolutionary history of a clade [80,81], which may decrease the power of comparative analyses, and lead to high probability of type I error [81]. In our case, the presence of at least one weapon is concentrated in two clades composed of species that predominantly spawn in exposed sites (figure 2). In both clades, each of the weapons investigated here is gained and *lost* frequently, an unexpected finding that does not fit our prediction. Thus, even though the small number of evolutionary transitions in the Leptodactylinae phylogeny might have increased the chances of type I error in our analysis, the results obtained from the HRMM do not align with our expectations. What the results show is that individual weapons are evolutionarily labile and that the dynamics of gains and losses are influenced by factors beyond the breeding site.

Evolutionary lability may stem from how much population-level effects influence each weapon over evolutionary time. For instance, in the myobatrachid frog *Crinia georgiana*, a species that spawn in exposed sites, the occurrence of hypertrophied arms used in male-male contests varies among populations, correlating with male density. In small to medium density populations, males possess hypertrophied arms due to intense pre-mating intrasexual competition, facilitating the profitable defense of reproductive resources. Conversely, in high male density populations, where pos-mating intrasexual competition is intense, males lack hypertrophied arms. In such populations, the mating system is a scramble competition, and males invest in larger testes rather than weapons [21,82]. Similarly, in a South African dung beetle community, male density was found to predict horn presence. Species lacking horns tended to exhibit high male density, whereas those with horns tended to exhibit low male density [83]. These two examples illustrate how, both within and between species, the presence of weapons can be influenced by male density. It remains unclear whether variations in male density among species spawning in exposed sites may influence the gain or loss of weapons in Leptodactylinae frogs. However, it appears that explosive breeding―characterized by large reproductive aggregations with high male density [5]―is more prevalent in frog species that spawn in exposed sites [84]. If this pattern holds true for the Leptodactylinae (e.g., [85]), it may explain why gains and losses of hypertrophied arms are more frequent in species that use this type of breeding site.

Contrary to hypertrophied arms, we found no report in the literature showing intraspecific variation in the presence of spines in frogs. The documented cases of variation are all interspecific and come from modifications in the shape of the thumb spine or attachment of muscles on the base of the thumb spine [82,83]. Also contrary to hypertrophied arms, which are labile weapons that appear and disappear frequently along evolutionary time of the Leptodactylinae, thumb spines are more stable across the evolution of the clade (figure 2). It appears, therefore, that spines exhibit higher levels of canalization within species, and we suggest that this canalization constrains evolvability, which represents a key determinant of evolutionary lability [86]. Given that thumb spines occur in other anuran clades [5,13], future studies could investigate whether the phylogenetic inertia of this trait is also higher than other weapons.

## 5. Conclusion

Our results underscore the significance of the type of breeding site in the evolution of weapons in a clade of Neotropical frogs. Through the application of two phylogenetic comparative methods employing complementary approaches, we argue that the consistent association between the presence of weapons and reproduction in exposed breeding sites, coupled with the enduring association between the absence of weapons and reproduction in concealed breeding sites, suggest a macroevolutionary interdependence between male weapons and type of breeding sites in the Leptodactylinae. In addition to the enduring association between the absence of weapons and reproduction in concealed breeding sites, our results also suggest a previously unexplored role of female mate choice in mediating evolution of male weapons. In species in which males do not force amplexus and female mate choice is primarily based on acoustic signals emitted by the males, direct male-male contests are rare, and weapons are mostly absent. Conversely, in species in which males force amplexus and female mate choice is limited, direct male-male contests for access to females are intense and weapons are mostly present. Finally, despite the evolutionary correlation between the presence or absence of weapons and type of breeding site, the dynamics of weaponry evolution is more intricate than anticipated. We found that both gains and *losses* of weapons occurred more frequently in species spawning in exposed sites compared to species spawning in concealed sites. We proposed some explanations for this unexpected pattern, aspiring to stimulate further investigations into the role of breeding site exposition on weaponry evolution not only in Leptodactylinae frogs, but also in other clades of anurans and external fertilizers, such as fish.

## Supporting information

Final dataset

Ultrametric tree

Methods details and additional results

Final dataset references

Genbank accession codes

