## Supplementary figures and images for "The evolution of male weapons is associated with the type of breeding site in a clade of Neotropical frogs"

### Ultrametric tree

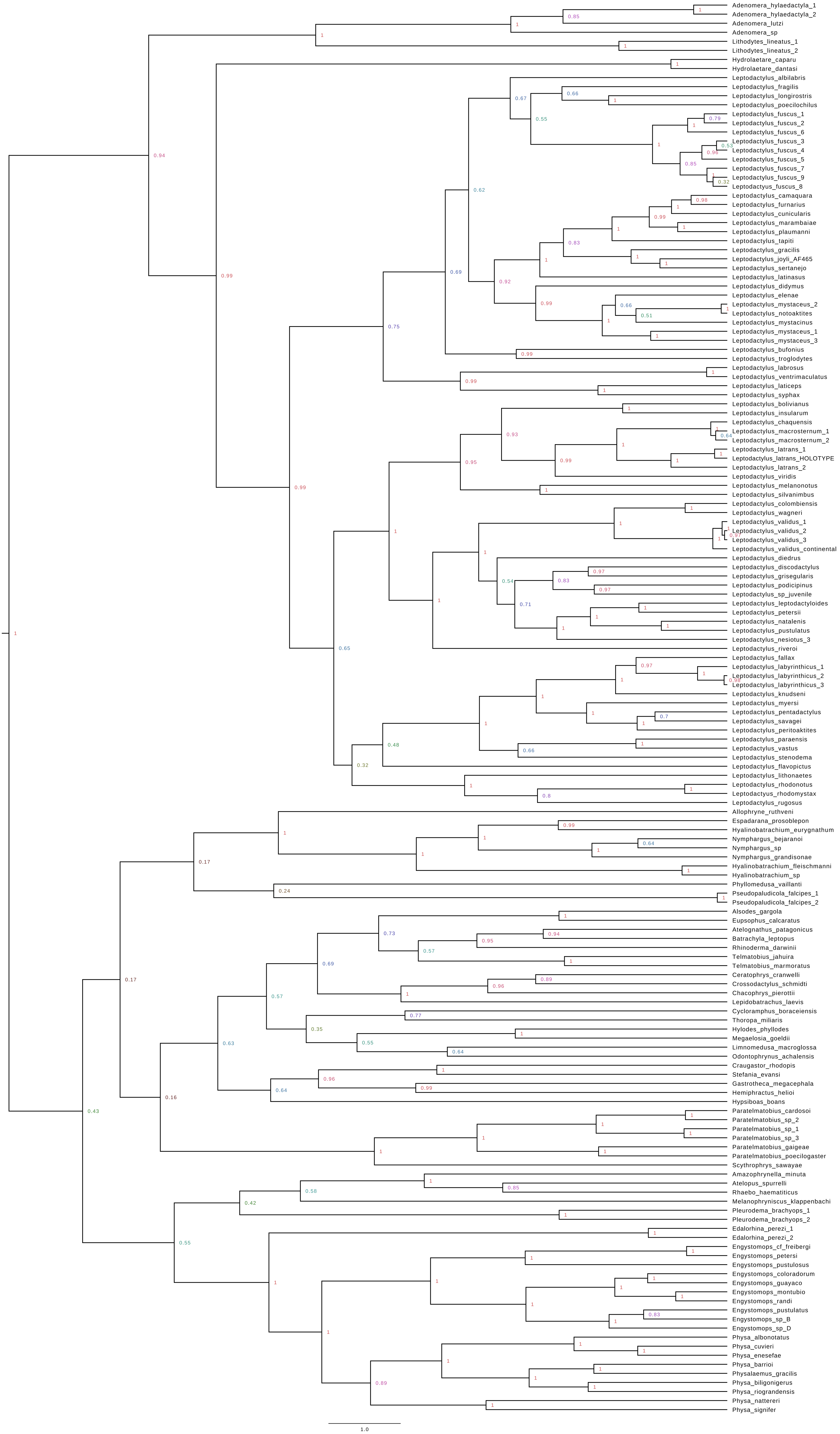
