## Supplementary material for "The evolution of male weapons is associated with the type of breeding site in a clade of Neotropical frogs": Methods details and additional results

**Supplementary Information S3**

**Material and Methods**

*(a) Tree construction*

For some Leptodactylinae species there was no molecular information available on GenBank, so we could not infer a larger tree based solely on molecular data. To circumvent this issue, we added the 10 species according to their putative sister species relationships along the midpoint of the branches using the ‘bind.tip’ function in the R package “phytools”. Table S1 below lists the species added and their sister pair relationships with species present in the original molecular phylogeny.

Table S1: Relationships between species added and sister species already present in the tree.

| Species added | Sister species relationship | Reference |
| --- | --- | --- |
| *Leptodactylus brevipes* | *Leptodactylus podicipinus* | Gazoni et al 2021 |
| *Leptodactylus spixi* | *Leptodactylus notoaktites* | Alves da Silva et al 2020 |
| *Leptodactylus cunicularius* | *Leptodactylus furnarius* | Carvalho et al 2020 |
| *Leptodactylus guianensis* | *Leptodactylus bolivianus* | Magalhães et al 2020 |
| *Leptodactylus paranaru* | *Leptodactylus latrans* *+* *Leptodactylus luctator* + *Leptodactylus payaya* | Magalhães et al 2020 |
| *Leptodactylus luctator* | *Leptodactylus latrans* + *Leptodactylus paranaru* + *Leptodactylus payaya* | Magalhães et al 2020 |
| *Leptodactylus payaya* | *Leptodactylus latrans* + *Leptodactylus luctator* + *Leptodactylus paranaru* | Magalhães et al 2020 |
| *Adenomera diptyx* | *Adenomera hylaedactyla 1* | Fouquet et al 2013 |
| *Adenomera andreae* | all other *"Adenomera"* | Fouquet et al 2013 |

*(b) D-test: using posterior predictive simulations*

Huelsenbeck and colleagues (2003) D-test uses posterior predictive simulations to compute a p value of the evolutionary association among traits. To do that, first we estimate the rates of transition between the states of the predictor trait (type of breeding site; binary) and the response trait (male weaponry; three states) independently (i.e., one separate Markov model for each trait). Then we use stochastic mapping simulations to estimate the distribution of ancestral states along the branches of the tree for the predictor and response traits (independently). Next, we compare the distribution of character mappings for both traits and compute the amount of evolutionary co-occurrence between each possible pair of states between the predictor and response traits.

The null distribution is generated by simulating new datasets using the same parameters estimated with the empirical data. This means that the null distribution has the same model complexity as the model fitted to the empirical data. After simulating new datasets (independently for the predictor and response traits), we estimate the same model and use the transition matrices to produce stochastic mappings and compute the amount of evolutionary association between the predictor and response traits. This process of simulating data based on empirical estimations and producing a (posterior) distribution of summary statistics is called *posterior predictive simulation* (or test). The resulting distribution is a null model that describes the expected amount of evolutionary association if the response and predictor traits had truly evolved independently. Finally, we use this null distribution to compute the p value for the D-test. Under Huelsenbeck et al. (2003) test of evolutionary correlation, traits are considered correlated if they co-occur across the branches of the tree more often than expected under the null model, which has the same complexity that alternative models.

**Results**

***(a)*** *Comparison with previous phylogenetic inference*

The relationships between *Adenomera* species present in both trees were similar. *Leptodactylus fragilis* is sister to all other *Leptodactylus* species in de Sá et al. (2014), but is more apical and closely related to *L. albilabris*, *L. longirostris*, *L. poecilochilus*, and *L. fuscus* in our tree. The relationship of these four species is similar in both trees. *Leptodactylus spixi* is absent in de Sá et al. (2014) tree and present in ours, closely related to *L. mystacinus*, *L. notoaktites* and *L. mystaceus* 2. The relationships between species of the *L. fuscus* species group are similar in both trees. In de Sá et al. (2014) *L. melanonotus* branches out from the node descendant from the MRCA of *Leptodactylus* and is closely related to *Hydrolaetare*. In our tree, however, *L. melanonotus* is more apical and closely related to the other species of the *L. latrans* species group; the other species of *L. melanonotus* species group present the same relationships in both trees. Species of the *L. pentadactylus* species group present the most similar relationships between the de Sá et al. (2014) tree and our tree (figure 4 of de Sá et al. 2014, figure 2 of this work).
